# Radiofrequency ablation remodels the tumor microenvironment and promotes systemic immunomodulation in pancreatic cancer

**DOI:** 10.1101/2022.01.07.475451

**Authors:** Erika Y. Faraoni, Nirav C. Thosani, Baylee O’Brien, Lincoln N. Strickland, Victoria Mota, Jarod Chaney, Putao Cen, Julie Rowe, Jessica Cardenas, Kyle L. Poulsen, Curtis J. Wray, Jennifer Bailey-Lundberg

## Abstract

**Background and Aims:** Pancreatic ductal adenocarcinoma (PDAC) is characterized by resistance to therapy. A major contributing factor to therapeutic failure is profound desmoplasia and a well-documented hypoxic tumor microenvironment (TME). In PDAC, several therapeutic approaches, including chemotherapy and radiation alone or combined with immune checkpoint inhibitors, have shown minimal therapeutic success, placing an imperative need for the discovery and application of innovative treatments. Endoscopic ultrasound guided radiofrequency ablation (EUS-RFA) is a promising immunomodulator therapy for PDAC. In this work, we *hypothesized* RFA promotes local and systemic stromal and immunomodulating effects that can be identified for new combination therapeutic strategies.

**Methods:** To test our hypothesis, a syngeneic PDAC mouse model was performed by symmetrically injecting 100k murine KPC cells in bilateral flanks of C57BL/6 female mice. RFA treatment initiated when tumors reached 200-500 mm^3^ and was performed only in the right flank. The left flank tumor (non-RFA contralateral side) was used as a paired control for further analysis.

**Results:** RFA promoted a significant reduction in tumor growth rate 4 days after treatment in RFA treated and non-RFA side contralateral tumors from treated mice when compared to controls. Histological analysis revealed a significant increase in expression of cleaved Caspase3 in RFA treated tumors. In addition, collagen deposition and CD31+ cells were significantly elevated in RFA side and non-RFA contralateral tumors from RFA treated mice. Proteome profiling showed changes in C5a and IL-23 in RFA responsive tumors, indicating a role of RFA in modulating intratumoral inflammatory responses.

**Conclusions:** These data indicate RFA promotes local and systemic anti-tumor responses in a syngeneic mouse model of PDAC implicating RFA treatment for local tumors as well as metastatic disease.

**Graphical Abstract:** 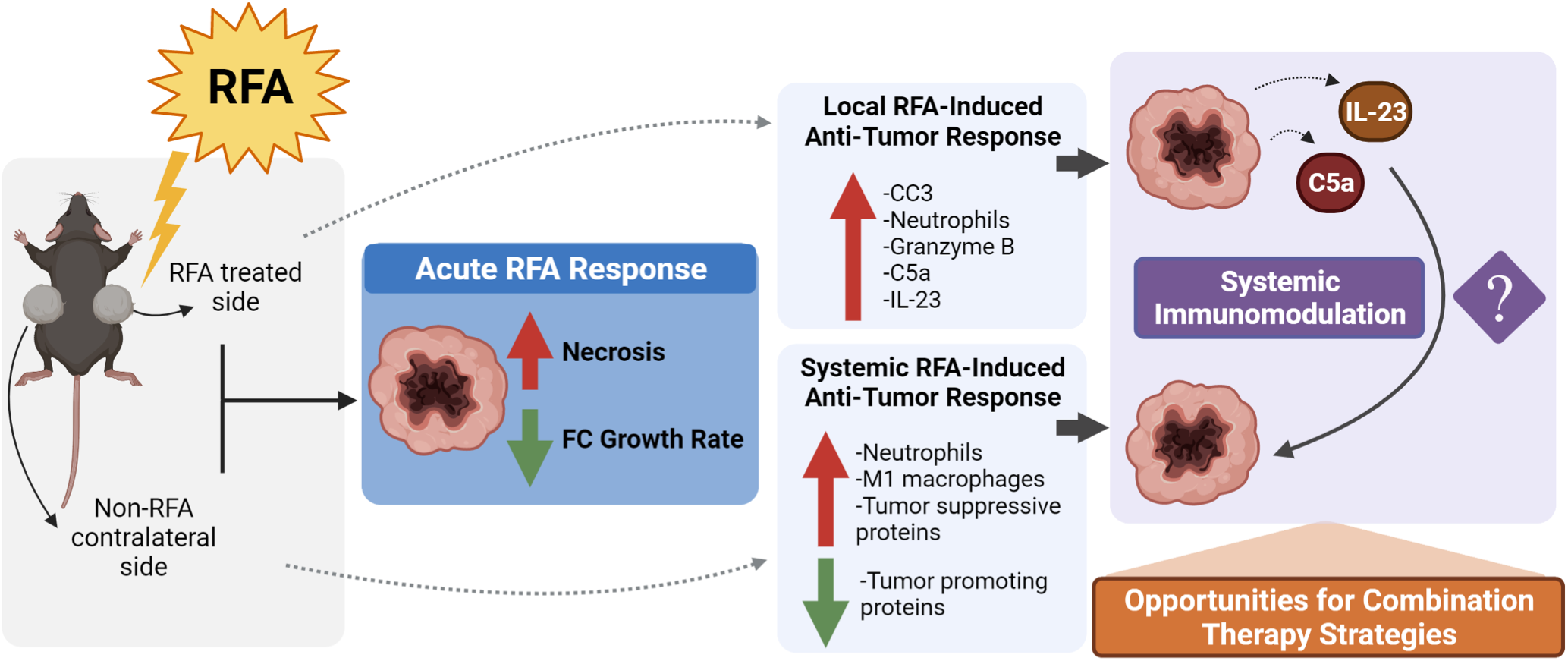

## INTRODUCTION

PDAC is projected to become the second deadliest cancer by 2025 ^1^ and is characterized by a prominent desmoplasia, which consists of cancer-associated fibroblasts (CAFs) and immune cells, whose dynamic nature play a crucial role in the development of PDAC ^2–5^ In pancreatic tumors, different CAFs and immune cell populations coexist in the TME with wide functional diversity, promoting both suppressive and supportive effects on tumor growth and playing a critical role in tumor development and therapy response ^6–8^ Numerous studies describe the complex and heterogeneous immune landscape of PDAC which includes the recruitment of immune cells that secrete immunosuppressive factors, such as chemokines (CXC12), cytokines (IL-1, IL-6, IL-10, TGFβ, TNF-α and GM-CSF), as well as the expression of cell-surface proteins, such as PD-L1 and CTLA4 which are checkpoint inhibitor molecules that confer inhibitory signals to the immune compartment and contribute to immune evasion^9^. In some PDAC patients, an increase in tumor associated macrophages (TAM) and a decrease in CD4+ and/or CD8+ T cells has been correlated with poor prognosis ^9,10^ indicating immunotherapeutic strategies to increase anti-tumor immunity can improve prognosis. Several therapeutic approaches, including chemotherapy and radiation alone or combined with immune checkpoint inhibitors have shown little success in PDAC, placing an imperative need for the discovery and application of innovative techniques ^11,12^.

Endoscopic ultrasound guided radiofrequency ablation (EUS-RFA) has the advantage of being minimally invasive and safe and is one of the newer FDA-approved techniques currently available for the treatment of a number of gastrointestinal malignancies including pancreatic cancer ^13–15^. Studies in both animal models and cancer patients show that RFA induces not only local burning/disruption of the tumor by heat but also generates coagulative necrosis in local RFA treated tumors which releases large amounts of cellular debris that represent a source of tumor antigens that can trigger a host adaptive immune response against the tumor ^16–18^. We have recently evaluated long-term outcomes of EUS-RFA in patients with advanced PDAC. EUS-RFA could be combined with standard of care chemotherapy in our patient cohort, was well tolerated and improved survival outcomes ^19^. Our clinical data indicate RFA in combination with standard of care chemotherapy is a promising therapeutic strategy. Other RFA preclinical and clinical studies have been done in treating advanced cancers at the metastatic site. In colorectal cancer liver metastases, RFA treatment of liver metastasis was shown to promote local and systemic Th1 type immune responses in both animal models and human patients. In these studies, RFA treatment played a significant role in inhibiting tumor recurrence from residual micro metastases or circulating tumor cells ^20,21^.

In PDAC, one study has been reported in mice using bilateral injections of Pan02 cells ^17^. This group reported decreased numbers of T regulatory cells, tumor-associated macrophages and tumor-associated neutrophils. Additionally, this study showed increased numbers of functional DCs, CD4+, and CD8+ T cells on day 3 after RFA treatment; however, the transient immune responses in this work lacked the ability to suppress tumor growth. In contrast, our preliminary preclinical work involves a syngeneic mouse model of PDAC, with bilateral injections of KPC cells. Unlike Pan02 cells, KPC cells express an oncogenic *Kras^G12D^* mutation present in the parental tumor allowing allografted tumors, either subcutaneously or orthotopically, retain a variable degree of desmoplasia, which better resembles the histological appearance and leukocyte complexity of the spontaneous parental tumor in KPC mice ^22^. Our results show a significant reduction in tumor growth rate 4 days after RFA treatment in RFA and non-RFA side contralateral tumors in treated mice compared to control tumors. This reduction in size was accompanied by significant upregulation of cleaved Caspase3 expression in RFA treated tumors and significant remodeling of the stroma. To better characterize the systemic response, Imaging Mass cytometry was done on non-RFA contralateral tumors and neighborhood analysis was used to observe spacial localization of immune cell types with epithelium. Surprisingly, while we quantified a significant increase in granzyme B in RFA treated tumors compared to controls, we did not observe a significant increase in granzyme B in non-RFA contralateral tumors. These results led us to *hypothesize* RFA initially promotes a strong local and systemic anti-tumor response, but tumor microenvironment remodeling and immunosuppressive mechanisms may also be elevated, which may compromise the long-term effects post ablation.

Our results are timely as therapeutic strategies to treat pancreatic cancer patients with locally advanced or metastatic disease remain limited. Current chemotherapy regimens such as modified FOLFIRINOX or gemcitabine plus nab-paclitaxel have shown survival advantage over gemcitabine alone and are the standard systemic treatment options for PDAC ^23^. Immunotherapies are gaining significant traction and recently a combination of TIGIT/PD-1 co-blockade in combination with CD40 agonism elicited anti-tumor responses in pre-clinical studies ^24^.

## RESULTS

### Radiofrequency ablation reduces PDAC tumor growth rate *in vivo*

To determine the effects of RFA *in vivo*, we established a syngeneic mouse model of pancreatic cancer and recorded tumor measurement every 4 days. After tumor ablation in one flank, we bilaterally monitored tumor size for another 4 days and compared tumor growth to a Sham control group which received no treatment (**Figure 1A**). RFA ablation did not affect body weight (**Figure S1A**). Comprehensive assessment of fold change (FC) in tumor growth rate 4 days before and after RFA treatment revealed a successful reduction with overall response in 14 mice (82.35%), from which more than 6 of them (42%) were categorized as bilateral responses including both RFA side and non-RFA side contralateral tumors (**Figure S1B**, see details in **Table 1**). Only three mice (17.65%) showed no acute response to RFA treatment on either side. In these three mice, the RFA treated tumors did not display any signs of necrosis indicating ablation, suggesting the absence of response in these three mice is due to technical failure (**Figure S2A**).

**Figure 1.**
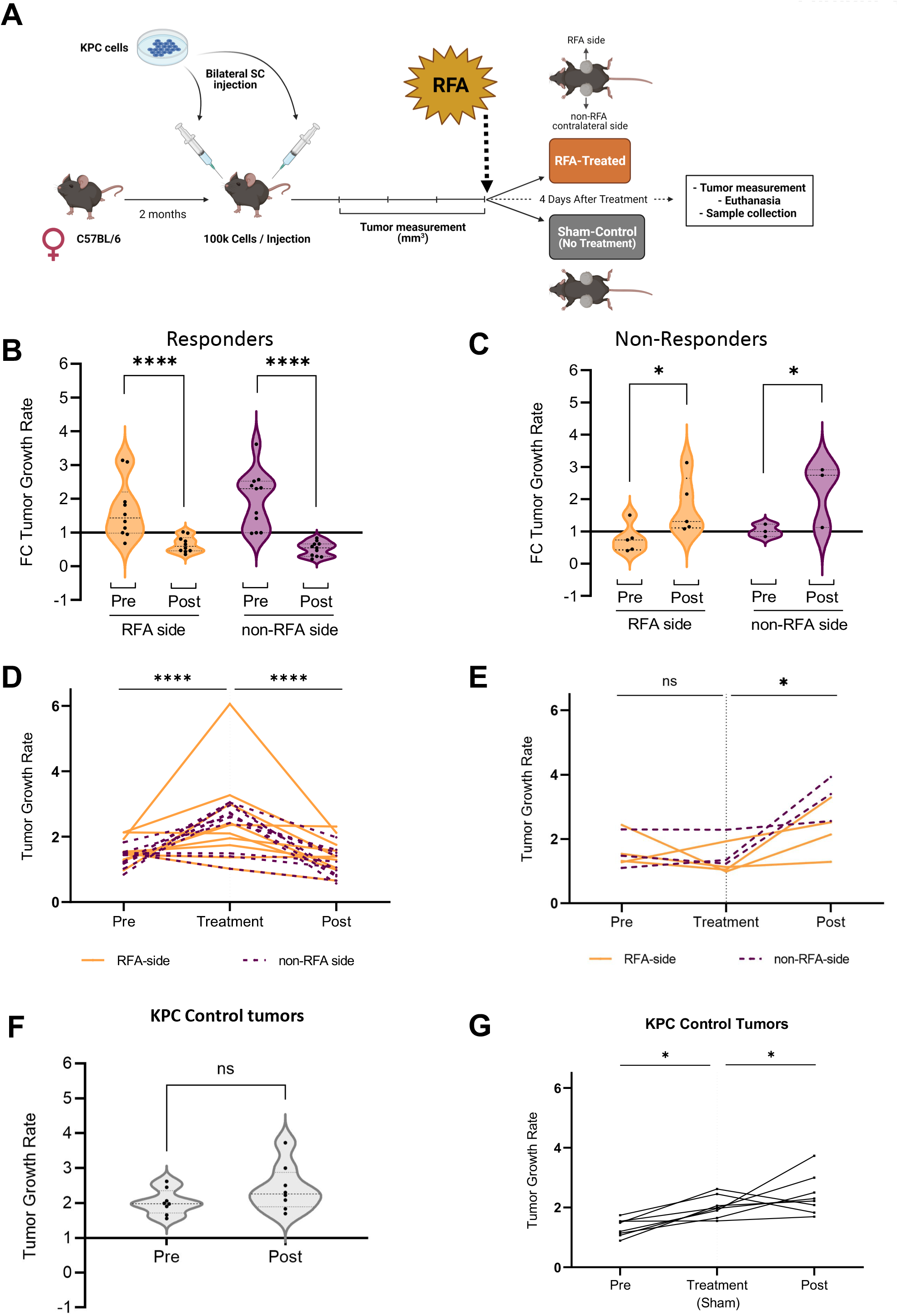
Radiofrequency ablation reduces PDAC tumor growth rate *in vivo*. **(A)** Experimental design of RFA treatment. To determine if RFA reduces tumor growth, we performed a bilateral injection model of 100k murine KPC cells. Tumor growth was measured semi-weekly and RFA was performed on one side when the tumors reached 500mm^3^. Tumor growth was monitored for another 4 days before mice were euthanized. **(B)** Decreased fold change (FC) in tumor growth rate 4 days pre and post RFA treatment was observed in RFA side (*n*=10) and non-RFA side (*n*=10) responsive tumors (p<0.0001). **(C)** Increased fold change (FC) in tumor growth rate 4 days pre and post RFA treatment was observed in RFA side (*n*=4) and non-RFA side (*n*=4) non-responsive tumors (p<0.05). **(D)** Response in RFA side (*n*=10) and non-RFA side (*n*=10) tumors presented a biphasic growth curve with increased (p<0.0001) and decreased (p<0.0001) rate pre and post RFA treatment. **(E)** Non-Responsive RFA side (*n*=4) and non-RFA side (n=4) tumors showed a significant increase in tumor growth rate 4 days pre and post RFA treatment (p<0.05) **(F)** At time of euthanasia, comprehensive assessment of fold change (FC) in tumor growth rate 4 days pre and post RFA treatment revealed no differences in control tumors (n=8; P=0.63). **(G)** Growth rate in KPC control group. Control sham treated tumors presented a continuous growth rate curve during the whole study (*n*=8; *P<0.05*). Data was analyzed by paired 2-way ANOVA to determine statistical significance. Interaction (p<0.0001).

**Table 1.**
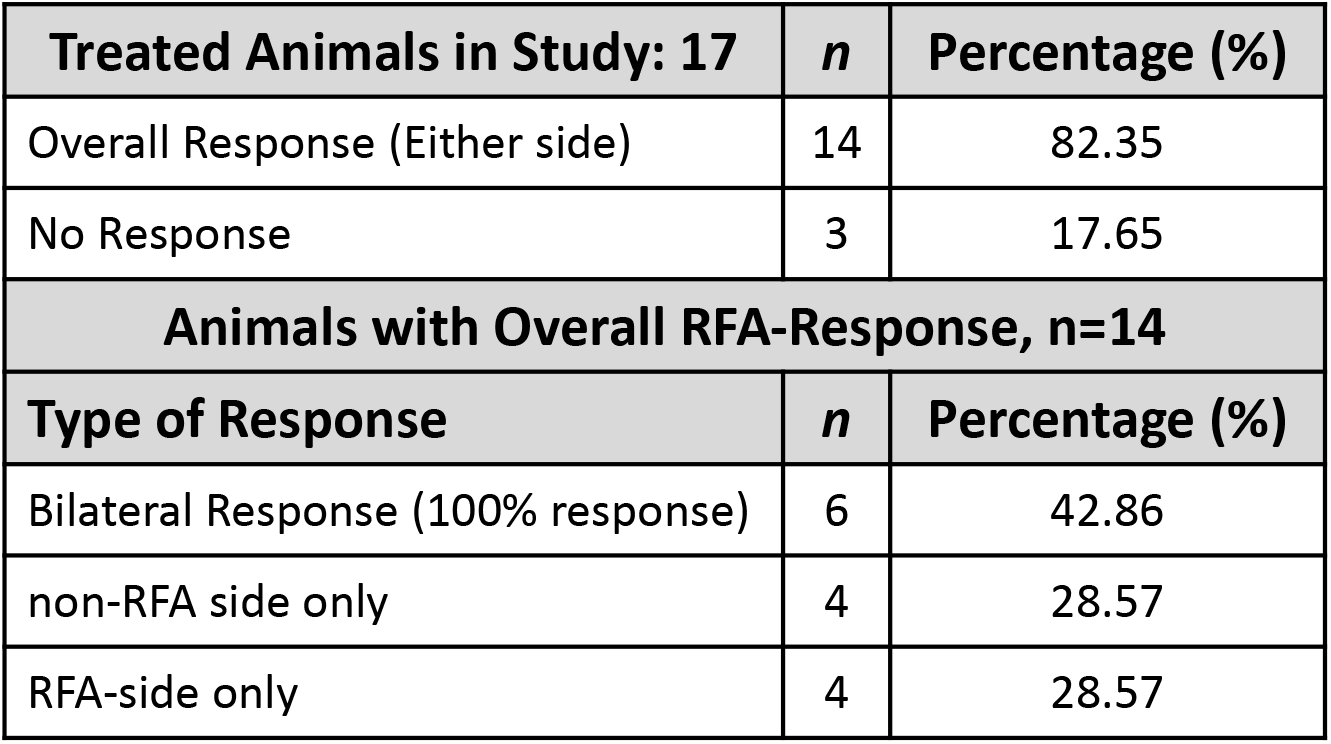
Animals treated with radiofrequency ablation (RFA) and type of response.

| **Treated Animals in Study: 17** | ***n*** | **Percentage (%)** |
| --- | --- | --- |
| Overall Response (Either side) | 14 | 82.35 |
| No Response | 3 | 17.65 |
| **Animals with Overall RFA-Response, n=14** |  |  |
| **Type of Response** | ***n*** | **Percentage (%)** |
| Bilateral Response (100% response) | 6 | 42.86 |
| non-RFA side only | 4 | 28.57 |
| RFA-side only | 4 | 28.57 |

In RFA responsive tumors, successful RFA ablation abruptly induced a three FC reduction in tumor growth rate 4 days after (Post) treatment in both the local (RFA side) and systemic (non-RFA) contralateral sides (**Figure 1B,** p<0.0001), while no difference was observed in control Sham-treated tumors (**Figure 1F,** p=0.63). Unexpectedly, in this group of mice, we observe a small subset of tumors (n=8) did not respond post RFA. These RFA non-responsive tumors doubled their FC in tumor growth rate 4 days after RFA treatment (**Figure 1C**, p<0.01), irrespective of their location.

To better comprehend the RFA ablation effect, we analyzed the growth rate curves in both RFA and non-RFA responsive tumors. Surprisingly, RFA responsive tumors irrespective of their location, presented a biphasic tumor growth rate curve with increased (p<0.0001) and decreased (p<0.0001) growth rate before (Pre) and after (Post) RFA treatment (**Figure 1D**). Like KPC control tumors, which continued to grow at a similar rate during the study (**Figures 1F** and **1G**), RFA treated non-responsive tumors presented a constant tumor growth rate before RFA treatment; however, they increased their tumor growth rate after ablation (**Figure 1E**, p<0.05).

### RFA promotes significant necrosis in both RFA treated and non-RFA tumors

To assess potential histopathological similarities in local and systemic responses between tumors 4 days after RFA treatment, we evaluated necrosis and apoptotic markers in responsive tumors. Composite images of hematoxylin and eosin staining revealed that both RFA side (p<0.05) and non-RFA contralateral side tumors (p<0.01) presented significantly increased necrotic area compared to control KPC tumors with no evident differences between sides (**Figures 2A** and **2D**). In non-responsive tumors, no differences were found in necrotic area between groups (**Figure S2B**).

**Figure 2.**
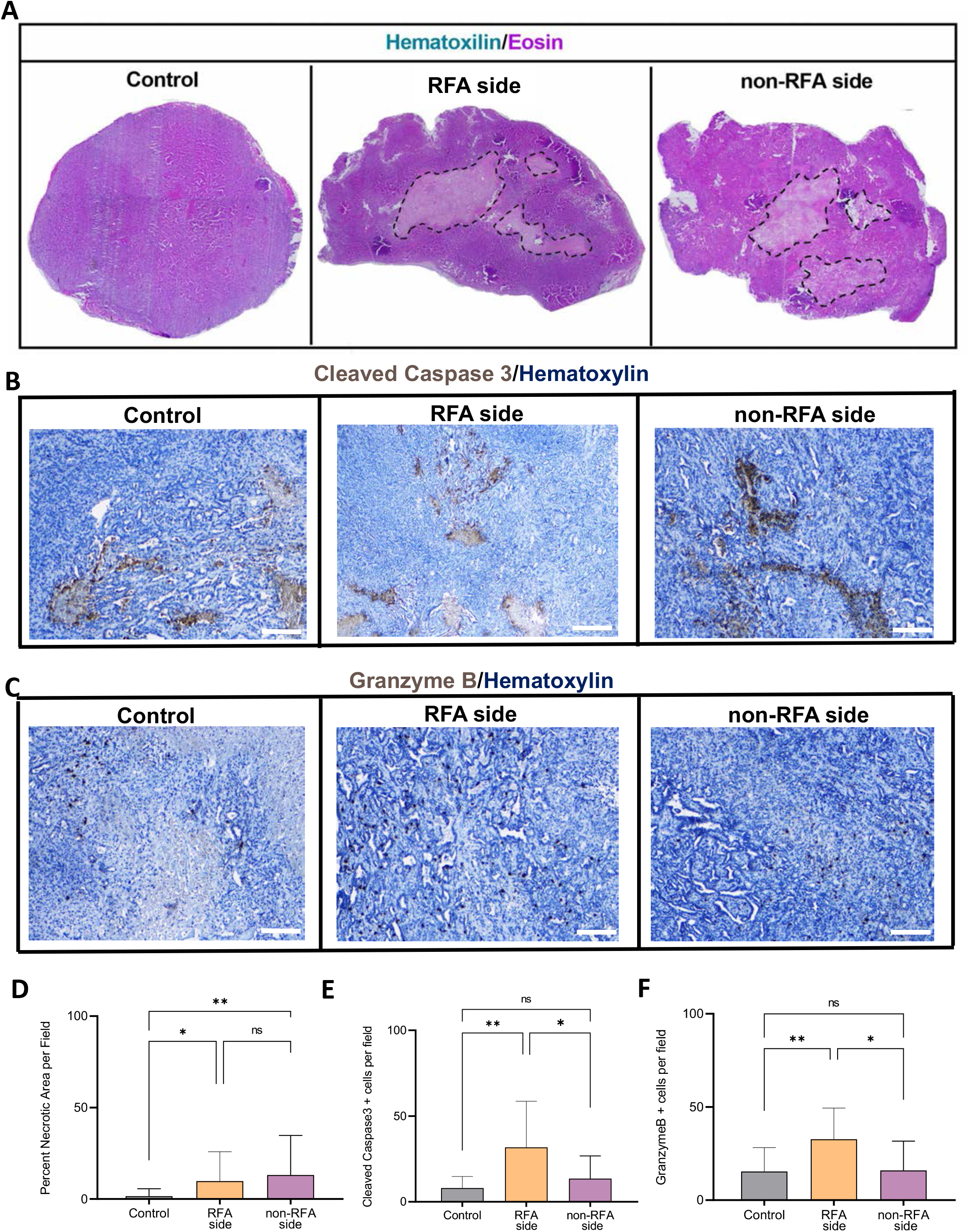
RFA increases necrotic area on both the RFA and non-RFA tumors compared to control KPC subcutaneous tumors. **(A)** Representative composite H&E staining of control, RFA, and non-RFA contralateral side tumors. **(B)** IHC staining for cleaved caspase 3 in control, RFA, and non-RFA side tumors. **(C)** IHC staining for granzyme B in control, RFA, and non-RFA contralateral side tumors. **(D)** ImageJ quantification of necrosis in control, RFA and non-RFA. RFA significantly increases necrosis on the RFA side (p<0.05; n=5) and non-RFA side tumors (p<0.01; n=7) compared to control tumors (n=5). **(E)** RFA increases cleaved caspase 3+ cells in the RFA side tumor compared to control (p<0.01) and non-RFA contralateral side tumors (p<0.05). **(F)** RFA significantly increases the number of granzyme B+ cells in the RFA side tumor compared to control (p<0.01) and non-RFA contralateral side tumors (p<0.05). Scale bars are 50uM. One-way ANOVA was used for group comparison.

Conversely, immunohistochemistry staining of cleaved caspase 3, which is considered a reliable marker for cells that are dying, or have died by apoptosis, revealed significantly increased staining only in the RFA treated side when compared to KPC control (p<0.05) and non-RFA contralateral side tumors (**Figures 2B** and **2E**).

Similarly, granzyme B staining was also significantly increased only in RFA treated tumors when compared to both KPC control (p<0.05) and non-RFA contralateral side tumors (**Figures 2C** and **2F**). These results suggest differential anti-tumor mechanisms of RFA response when comparing local and systemic anti-tumor immunity despite similar necrotic induction after RFA ablation.

### RFA local response promotes C5/C5a and IL-23 secretion and neutrophil infiltration

To determine the nature of RFA response in locally ablated tumors, we performed a membrane-based antibody array in tumor homogenates and found that RFA significantly increases C5/C5a and IL-23 expression in RFA locally ablated tumors compared to KPC control (**Figure 3A**, p<0.0001). Based on the capability of both molecules to stimulate neutrophil recruitment, we assessed neutrophil tumor abundance by immunohistochemistry and found it was significantly elevated in both RFA (p<0.0001) and non-RFA contralateral side tumors (p<0.001) compared to KPC control (**Figures 3B** and **3C**). Moreover, we also observed elevated myeloperoxidase (MPO) staining in areas with high density of neutrophils indicating a prominent anti-tumor neutrophil response after ablation in both RFA and non-RFA contralateral side tumors (**Figure 3D**).

**Figure 3.**
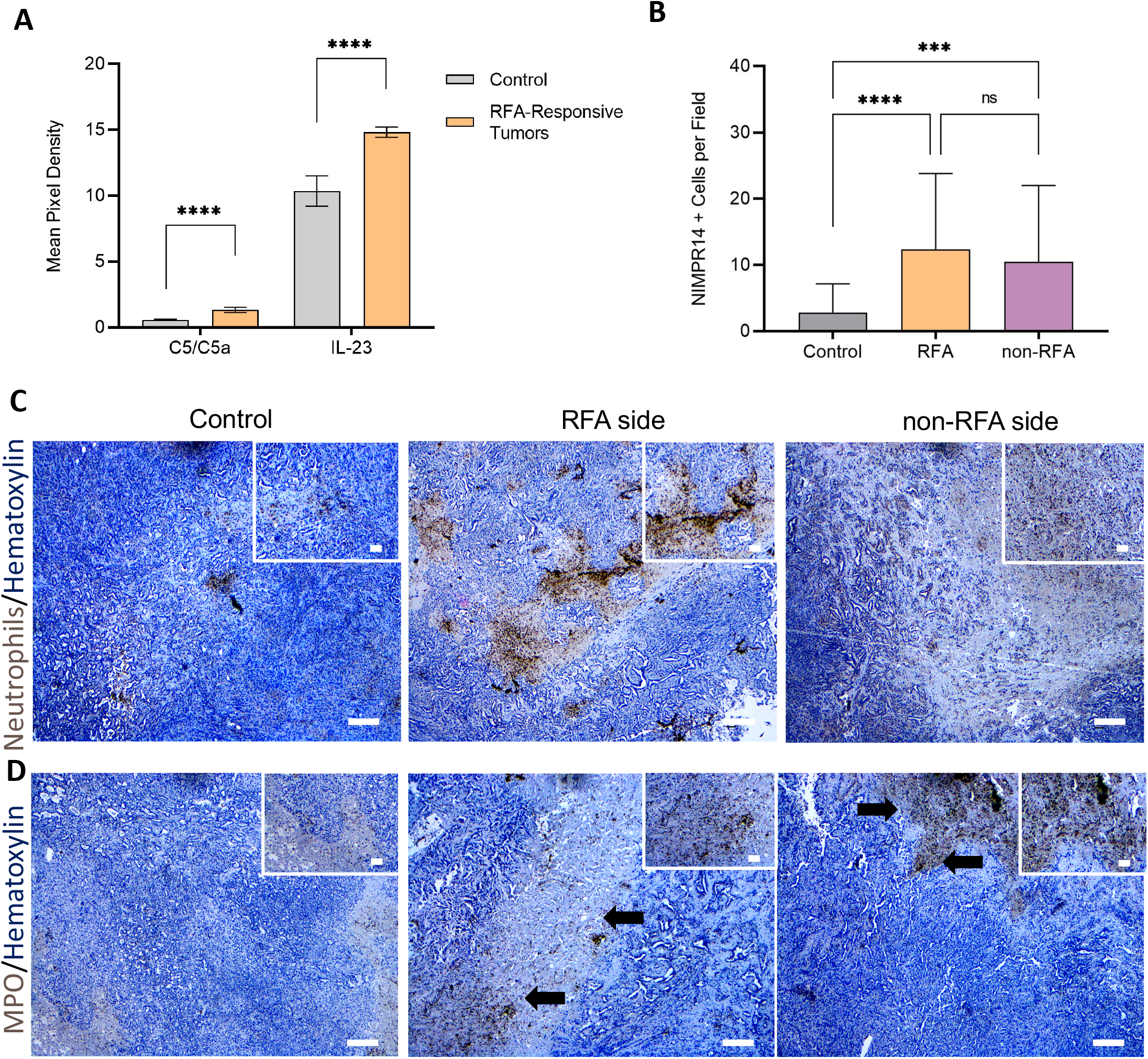
RFA local response increases C5/C5a and IL-23 secretion and neutrophil infiltration. **(A)** RFA significantly elevates C5 and IL-23 on the RFA side tumors (p<0.0001). An unpaired student’s t-test was used to evaluate significance. **(B)** We evaluated neutrophil abundance using IHC and quantified **(C)** number of neutrophils per field using ImageJ software. We observed a significant increase in neutrophils in RFA treated tumors (p<0.0001) and contralateral non-RFA side tumors (p<0.001) compared to control tumors. One-way Anova was used for group comparison. **(D)** Myeloperoxidase staining was used to study anti-tumor neutrophil abundance in RFA treated and contralateral tumors. We observed MPO staining in areas with high density of neutrophils (black arrows) indicating a prominent anti-tumor neutrophil response post RFA.

### RFA remodels the TME in local and non-RFA contralateral side tumors

As the immune system orchestrates key roles in neovascularization, collagen deposition, tissue remodeling and is important to characterize for response to immunotherapy, ^25–27^ we wanted to study the effects of RFA ablation on tumor fibrosis and angiogenesis. We performed trichrome staining in KPC control tumors and compared RFA side and non-RFA contralateral side tumors (**Figure 4A**). We quantified the percentage of collagen per field and observed both RFA (p<0.001) and non-RFA contralateral side (p<0.0001) tumors presented elevated levels when compared to KPC control tumors (**Figure 4C**). Similarly, both treated RFA (p<0.05) and non-RFA contralateral side tumors (p<0.05) presented increased CD31+ cell staining when compared to control samples (**Figures 4B** and **4D**), suggesting an active RFA-induced remodeling of the pancreatic cancer TME in both locally treated and distant non-treated sites.

**Figure 4.**
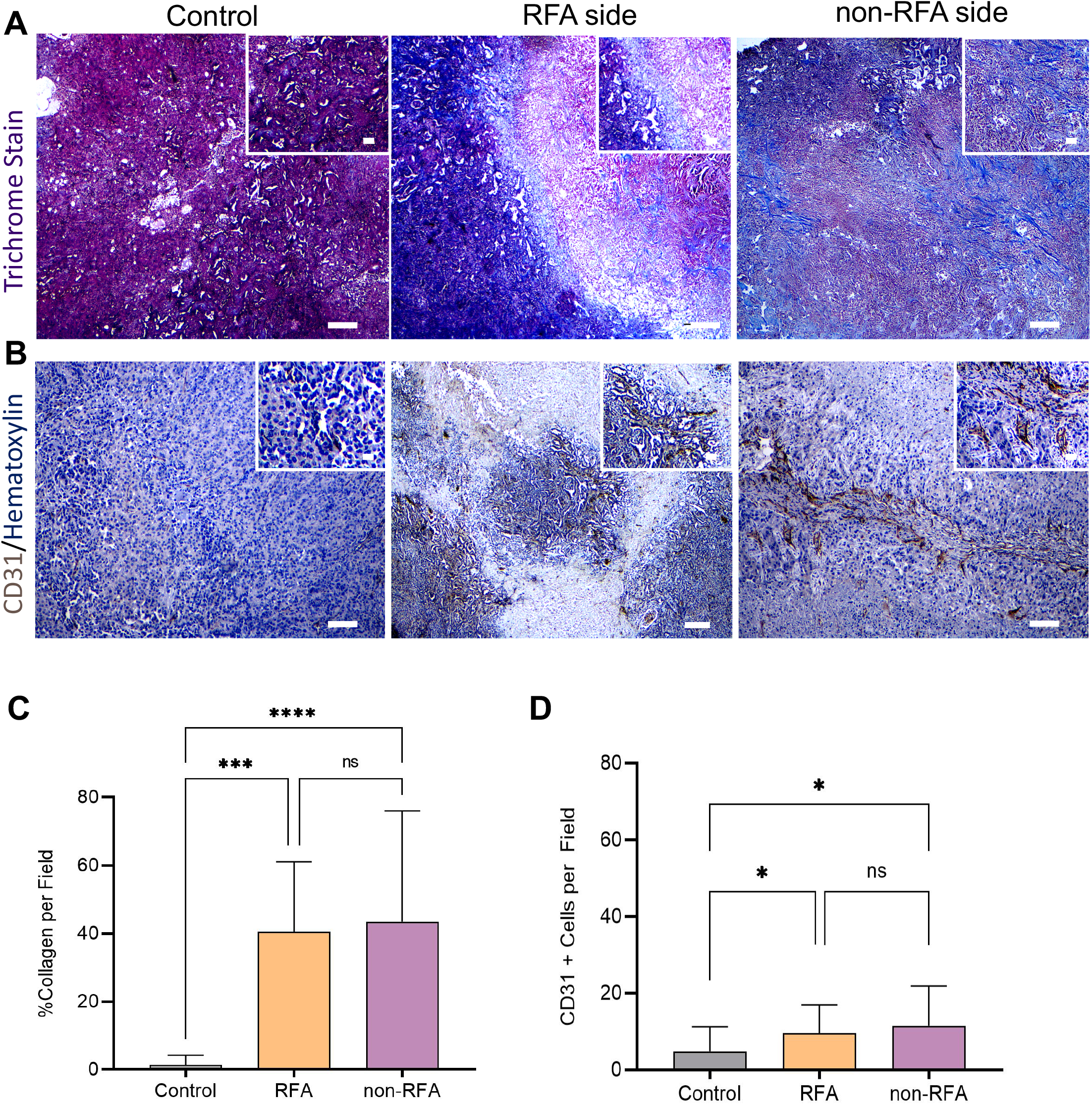
RFA induces significant collagen deposition in RFA and contralateral/non-RFA tumors. **(A)** Representative trichrome staining on control, RFA and non-RFA tumors. **(B)** Representative CD31 immunohistochemistry staining on control, RFA and non-RFA tumors. **(C)** ImageJ quantification of percent collagen+ tissue per field. RFA significantly increases collagen deposition in both the RFA treated (p<0.001) and non-RFA treated tumors (p<0.0001). One-way ANOVA was used for group comparison. **(D)** ImageJ quantification of percent collagen CD31 + cells per field. RFA significantly increases CD31 expression in both the RFA treated (p<0.05) and non-RFA treated tumors (p<0.05). Unpaired student’s t test in Prism Graphpad software was used to compare groups.

### RFA tumor microenvironment remodeling in human pancreatic tumors

To examine the impact of EUS-RFA in human PDAC, we used a hematoxylin and eosin stain on a resected PDAC specimen from a patient that received three ablation treatment sessions. The tumor was localized at the head of pancreas from a patient with locally advanced Stage III pancreatic cancer. The H&E image shows adjacent RFA-Ablated and Non-Ablated areas (**Figure 5A**). Limited epithelial cells were observed in the ablated region. Histopathological analysis of higher magnification RFA-ablated and non-ablated areas (**Figure 5B**, upper and lower panel, respectively) showed evident signs of necrotic tissue and collagen deposition as evaluated by H&E analysis and trichrome staining (**Figure 5B**, Middle panels) and increased angiogenesis as evidenced by CD31+ cells (**Figure 5B,** Right panels), with the latter more pronounced in the RFA ablated area compared to non-ablated region.

**Figure 5.**
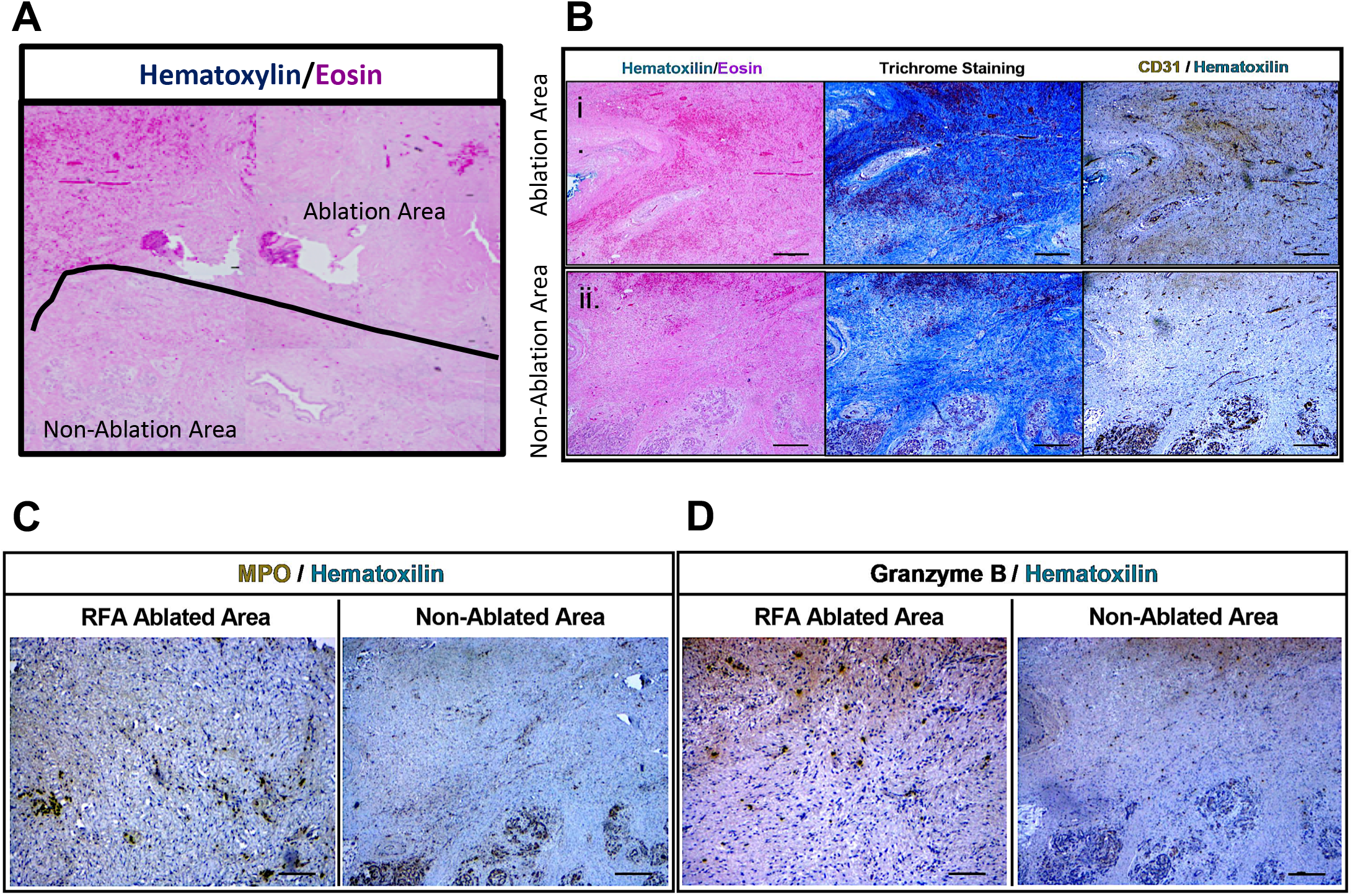
RFA ablation induces tumor microenvironment remodeling in human pancreatic tumors. **(A)** Representative composite H&E staining of human pancreatic cancer tumor showing both RFA-ablated and non-ablated areas. **(B)** H&E staining (Left) at 4X of human pancreatic cancer tumor in RFA-ablated (Upper panel Figure i.) and non-ablated (Lower panel Figure ii.) areas. Trichrome staining (Middle) at 4X of human pancreatic cancer tumor in RFA-ablated (Upper panel Figure i) and non-ablated (Lower panel Figure ii) areas. CD31 immunohistochemistry staining (Right) at 4X of human pancreatic cancer tumor in RFA-ablated (Upper panel Figure i) and non-ablated (Lower panel Figure ii) areas. **(C)** MPO immunohistochemistry staining of human pancreatic cancer tumor in RFA-ablated (Left 10X) and non-ablated (Right 4X) areas. **(D)** Granzyme B immunohistochemistry staining of human pancreatic cancer tumor in RFA-ablated (Left 10X) and non-ablated (Right 4X) areas. Scale bars are 50uM.

Based on the immune infiltrates observed in our preclinical mouse model, we also analyzed MPO and granzyme B by immunohistochemistry. Similar to what we found in murine subcutaneous tumors, we observed a significant presence of MPO staining suggesting active secretion of this inflammatory enzyme from immune cell infiltrates in both RFA-ablated and non-ablated areas (**Figure 5C**). Likewise, granzyme B was also present in both areas (**Figure 5D**) indicating a potential local activation of natural killer cells and cytotoxic T cells in human pancreatic tumors after RFA. These results are consistent with the significant induction of granzyme B + cells in our RFA treated murine tumors.

### RFA systemic response is associated with a significant induction of anti-tumor M1 macrophages, colocalizing with activated CD8+ T cells and anti-tumor neutrophils

A surprising result from our experiment was the potent acute reduction in FC tumor growth rate in the non-RFA contralateral side tumors. Notably, while we observed an increase in granzyme B + staining in these tumors, we did not quantify a significant induction of granzyme B in the non-RFA contralateral tumors. Thus, we wanted to comprehensively evaluate the immune cell composition of these tumors to define the cellularity of the immune cell microenvironment in tumors with systemic immunomodulation. To assess the abundance of immune cell infiltrates in distant non-treated tumors, we evaluated by Imaging Mass Cytometry (IMC) three regions from contralateral non-RFA side tumors (**Figure 6A**). Cluster identities are shown in **Figure 6B**. In analyzing the immune composition on the non-RFA contralateral side, IMC data quantification showed increased abundance of F480+CD44+ macrophages (**Figure 6A** and **6E**). tSNE plots were used to visualize cluster abundance (**Figure 6C**) and IMC Neighborhood analysis showed F480+CD44+ macrophages strongly colocalized with activated CD8+CD44+ T cells and anti-tumor neutrophils expressing Ly6G+CD11b+CD44+ (**Figures 6D** Clusters 19, 9 and 12).

**Figure 6.**
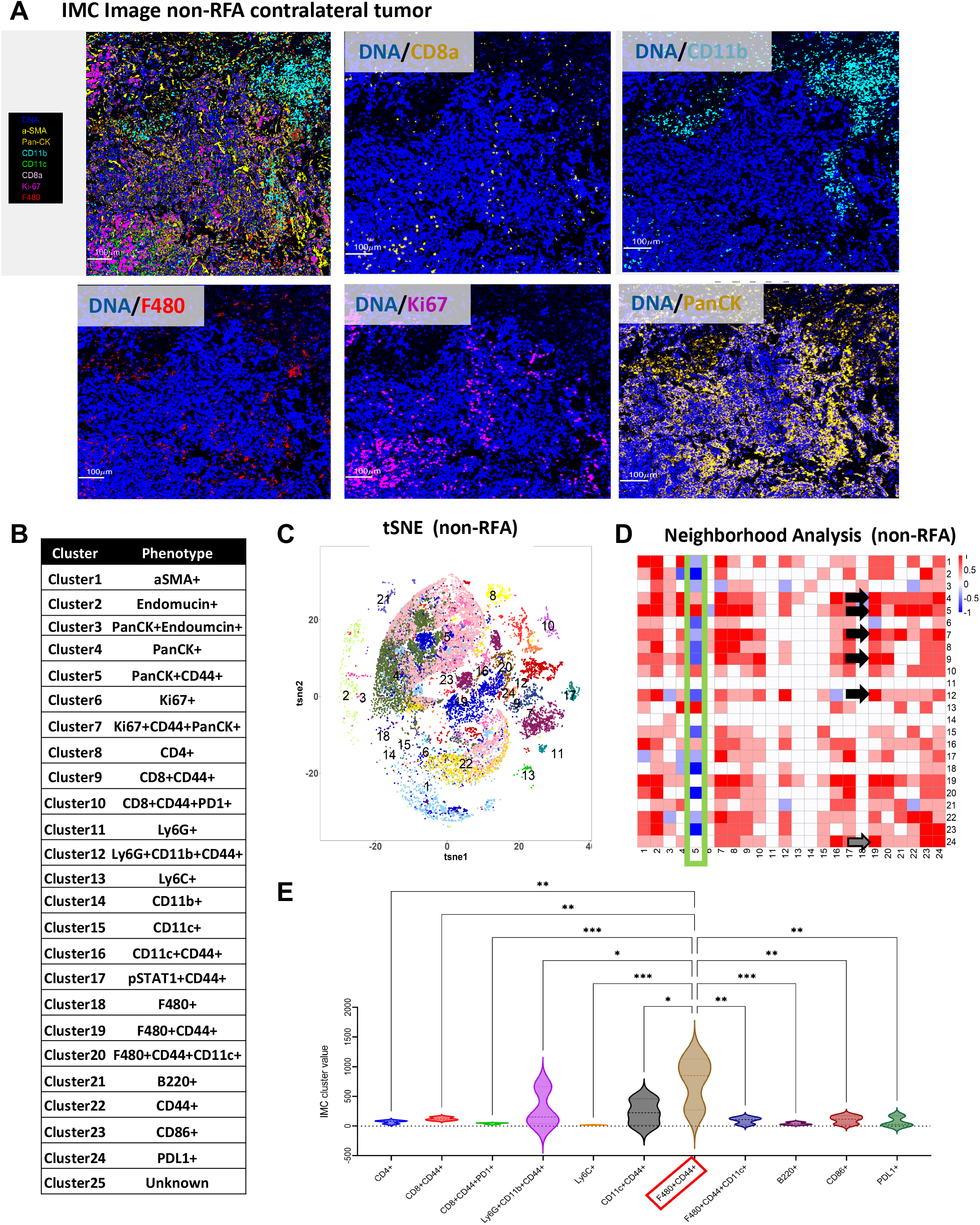
RFA systemic response is associated with a significant induction of anti-tumor M1 macrophages. To determine the abundance of immune cell infiltrates on the contralateral side tumors, we used Imaging Mass Cytometry (IMC) and analyzed three regions from the non-RFA treated contralateral side. **(A)** IMC fluorescent images of contralateral tumors. **(B)** Table with Cluster and Cell Phenotype information of Neighborhood analysis of IMC data. **(C)** tSNE plots of IMC clusters from non-RFA contralateral tumors. **(D)** Neighborhood analysis of IMC data indicating Cluster 5 with CD44+ cells (Green Box) is decreased is association with Cluster 6 (Ki67), Cluster 7 (CD4+), Cluster 8 (CD8+CD44+), Cluster 12 (Ly6G+CD11b+CD44+), Cluster 15 (CD11c+) and Cluster 23 (CD86+) cells. Similar patterns of association were observed in PanCK+ cells (cluster 4) between groups. Cluster 19 with F480+CD44+ Macrophages in non-RFA responding tumors colocalized with Clusters 4 and 5 (PanCK+ and PanCK+CD44+ cells, respectively), Cluster 7 (Ki67+CD44+PanCK+ cells), Cluster 9 (CD8+CD44+ cells), Cluster 12 (Ly6G+CD11b+CD44+ cells) and Cluster 24 (PDL1 + cells). **(E)** IMC data quantification reveals a significant induction of F4/80+CD44+ macrophages (red box). *p<0.05; **p<0.01; ***p<0.001. A One-way ANOVA was used for significance in Prism GraphPad Software.

Neighborhood analysis also evidenced a significantly decreased abundance of PanCK+CD44+ cells, a potential tumor stem cell population, adjacent to immune cell subtypes (**Figure 6B** -Cluster 5- and **Figure 6D** -Green Box-). In contrast, immune cell clusters were observed in close association to PanCK+ cells. A significant colocalization of PanCK expressing tumor cells, PanCK+CD44+ cells (Clusters 4 and 5, respectively) and proliferating cells expressing Ki67+CD44+PanCK+ (Cluster 7) was significantly associated with F480+CD44+ macrophages (**Figure 6D**, black arrows; Cluster 19) suggesting anti-tumor macrophages contribute significantly to the systemic response and increased necrosis in the non-RFA contralateral tumors. Notably, this study also revealed these macrophages colocalized with cells expressing PD-L1 (Cluster 24) indicating checkpoint blockade mechanisms may increase after the initial RFA antitumor response and combination anti-PD-L1/PD-1 with RFA may improve overall responses. Consistent with the neighborhood analysis, comprehensive IMC data quantification from these tumors revealed F480+CD44+ macrophages were significantly more prominent in these tumors compared to CD4+, CD8+CD44+, Ly6C+, CD11C+CD44+, F480+CD44+CD11c+, B220+ or CD86+ cells (**Figure 6E**).

### Large scale proteomics defines proteins altered in non-RFA contralateral tumors

In addition to histologic analysis, a large-scale functional proteomics platform using Reverse Phase Protein Array (RPPA) at MDAnderson Cancer Center was used to determine mechanisms altered in non-RFA tumors compared to control tumors. This functional proteomic approach characterizes protein expression and post translational modifications for over 400 proteins. Significantly downregulated proteins included pAkt^T308^, Cdc42, Gata6 and IGFRb (p<0.05), all important drivers of PDAC tumorigenesis (**Figure S3A**). Significantly increased proteins included INPP4b, MIG6, VHL and DDR1 (p<0.05). These proteins have tumor-suppressive functions in cancer and may be important determinants of the post-RFA systemic response (**Figure S3B**).

## METHODS

### Immunohistochemistry

Tissues were fixed in zinc buffered formalin, processed according to standard protocols, and embedded in paraffin. The unstained sections were baked at 60 °C for 45□minutes. The sections were deparaffinized with Histoclear and rehydrated. Antigen retrieval was performed using heat-mediated microwave methods and the antigen unmasking solution (Vector Laboratories, H-3300 pH 6) was used for cleaved caspase 3, NIMPR-14, and MPO antibodies. The antigen unmasking solution (Abcam 100X Tris-EDTA pH 9) was used for the CD31 and Granzyme B antibodies. All sections were blocked for one hour in 10% FBS in PBST and primary antibodies (**Table S1**) were incubated overnight at 4□°C. Secondary antibodies were used at 1:500 and incubated at room temperature for 30□minutes. The Vectastain ABC kit Peroxidase Standard (Vector Laboratories, PK4000) and DAB Peroxidase (HRP) Substrate kit (Vector Laboratories, SK-4100) were used. Slides were mounted in Cytoseal XYL (Epredia 8312-4) mounting media.

### Human Image

Human post resection image is deidentified to all research personnel and in compliance with UTHealth human subjects research.

### Trichrome Staining

The Abcam Trichrome Stain (ab150686) kit was used for connective tissue staining. Tissues were fixed in zinc buffered formalin, processed according to standard protocols, and embedded in paraffin. The sections were deparaffinized with Histoclear and rehydrated. Bouin’s Fluid was preheated to 60° C and sections were incubated for 60 minutes. Sections were then stained with Weigert’s Iron Hematoxylin for 5 minutes, Biebrich Scarlet for 15 minutes, Phosphomolybdic Acid for 12 minutes, Aniline Blue Solution for 20 minutes, and Acetic Acid Solution for 5 minutes. Sections were dehydrated with Histoclear. Sections were mounted in Cytoseal XYL (Epredia 8312-4) mounting media.

### ImageJ Methods

Area of necrosis on H&E sections was measured using ImageJ (http://imagej.nih.gov/ij/) software. All fields of pancreatic H&E composites were used for quantification. Necrosis was calculated as percentage of total pancreatic area. Freehand selection tool was used to isolate necrosis versus normal pancreatic tissues. Images for composite H&Es were taken in 4X. For collagen fiber quantification, Abcam trichrome-stained pancreatic sections were used. Color intensity was used to discern between true collagen staining and background staining. Freehand selection tool of ImageJ software was used to quantify the collagen area in five randomly chosen fields of each pancreatic section. Fibrosis was calculated as percentage of total pancreatic area. In total, 5 fields per slide were selected for quantification and all mice were analyzed per group. For IHC quantification, color intensity was used to discern between true IHC staining and background staining. Freehand selection tool of ImageJ software was used to quantify the IHC positive area in five randomly chosen fields of each pancreatic section. Positive staining was calculated as percentage of total pancreatic area. In total, 3-5 fields per slide were selected for quantification and all mice were analyzed per group.

### Imaging mass cytometry (IMC)

Metal-labeled antibodies were prepared according to the Fluidigm protocol. Antibodies were obtained in carrier/protein-free buffer and then prepared using the MaxPar antibody conjugation kit (Fluidigm). After determining the percent yield by absorbance measurement at 280 nm, the metal-labeled antibodies were diluted in Candor PBS Antibody Stabilization solution (Candor Bioscience) for long-term storage at 4°C. Antibodies used in this study are listed in Table E1 ^28^.

Tumor sections were baked at 60°C overnight, then dehydrated in xylene and rehydrated in a graded series of alcohol (ethanol absolute, ethanol: deionized water 90:10, 80:20, 70:30, 50:50, 0:100; 10 minutes each) for imaging mass cytometry. Heat-induced epitope retrieval was conducted in a water bath at 95°C in Tris buffer with Tween20 at pH 9 for 20 minutes. After immediate cooling for 20 minutes, the sections were blocked with 3% bovine serum albumin in tris-buffered saline (TBS) for 1 hour. For staining, the sections were incubated overnight at 4°C with an antibody master mix. Samples were then washed 4 times with TBS/0.1% Tween20. For nuclear staining, the sections were stained with Cell-ID Intercalator (Fluidigm) for 5 minutes and washed twice with TBS/0.1% Tween20. Slides were air-dried and stored at 4°C for ablation.

The sections were ablated with Hyperion (Fluidigm) for data acquisition. Imaging mass cytometry data were segmented by ilastik and CellProfiler. Histology topography cytometry analysis toolbox (HistoCAT) and R scripts were used to quantify cell number, generate tSNE plots, and perform neighborhood analysis. For all samples, tumor and cellular densities were averaged across 3 images per tumor, with n□=□3 per group.

### Animal Model

All mouse model procedures are in compliance with UTHealth’s CLAMC Animal Welfare Committee Review and approved on Dr. Bailey’s AWC protocol. 1X10^5^ KPC cells in PBS: Matrigel mix (1:1) were injected in the left and right flanks of female C57BL/6mice. Tumor size was calculated twice per week with vernier caliper. Tumor volume was calculated as (length × width × width)/2 in mm^3^. Radiofrequency ablation was performed on the right-side tumor once tumors were 200-500mm^3^. Tumor Growth Rate was calculated for each tumor pre and post RFA treatment as the ratio between 4-day consecutive tumor measurements (Ex: Growth rate Pre: Measurement 2 / Measurement 1; Growth rate Treatment: Measurement 4 / Measurement 3). Fold Change (FC) in tumor growth was calculated as the ratio between consecutive Growth Rate Measurements. (Ex: Fold Change (FC) Pre: Growth rate Treatment / Growth rate Pre; Fold Change (FC) Post: Growth rate Post / Growth rate Treatment).

### Tumor Proteome Array

The Proteome Profiler™ Array Mouse Cytokine Array Panel A (ARY006, R&D Systems, Minneapolis, MN, USA) was used to measure the protein expression levels of 40 cytokines and chemokines in KPC tumor tissues. Protein concentrations were quantitated using a total protein assay. The array was inoculated with 200 μg proteins, and samples were treated according to the product specification. Briefly, protein samples were diluted and mixed with a cocktail of biotinylated detection antibodies and then incubated with a mouse chemokine or cytokine array membrane. After washing, streptavidin-conjugated horseradish peroxidase and chemiluminescent detection reagents were added. Array images showing chemiluminescence signals were obtained using a ChemiDoc MP (Bio-Rad Laboratories) and analyzed by densitometry for integral optical density using Image Lab software. The optical density of each pair of chemokine or cytokine spots was normalized to the corresponding KPC control tumor spots.

### Radiofrequency ablation

RFA treatment was initiated when tumors reached 200-500 mm^3^. Mice were anesthetized with isoflurane via a Patterson Scientific Posi-Vac nose cone. A small incision was made in the right-side tumor to insert the Habib EUS RFA probe perpendicular to the skin in the center of the tumor. Average power delivered was 1.82 W over 10-20 seconds. The non-RFA side contralateral tumor did not receive RFA treatment. Mice were observed for signs of pain or discomfort post-ablation.

### Functional Proteomics/RPPA Analysis

Control and non-RFA contralateral side tumors were collected and lysed using lysis buffer (1% Triton X-100, 50mM HEPES, pH 7.4, 150mM NaCl, 1.5mM MgCl2, 1mMEGTA, 100 mM NaF, 10mM Na pyrophosphate, 1mM Na3VO4, 10% glycerol), containing freshly added protease and phosphatase inhibitors (Sigma Aldrich, St. Louis, MO) and protein concentration was measured by bicinchoninic acid (BCA) method.

Protein concentrations of samples were adjusted to 1mg/ml with lysis buffer. Cell lysates were serially diluted two-fold for 5 dilutions (from undiluted to 1:16 dilution) and arrayed on nitrocellulose-coated slides in an 11×11 format. Samples were probed with antibodies by tyramide-based signal amplification approach and visualized by DAB colorimetric reaction. The slides were analyzed and protein expression quantitated with the use of Array-Pro Analyzer. All the data points were normalized for protein loading and transformed to linear value, designated as “Normalized Linear”.

## DISCUSSION

In this study, we report local and systemic stromal modulation acutely induced by RFA. A number of preclinical mouse models have shown RFA is a potent mediator of antitumor responses as the ablation augments release of tumor antigens which elevates antigen-specific T cell responses ^15,17,29–31^. These effects have predominantly been studied in the local RFA ablation region. In this study, we observed an acute significant reduction in FC tumor growth rate in both the RFA and non-RFA contralateral subcutaneous tumors compared to control KPC tumors. In addition, we observed significant changes to the TME in both the RFA treated tumors and the non-RFA contralateral side tumors. These changes included a significant induction of necrosis and collagen expressing stroma as well as a significant increase in CD31+ vascular cells adjacent to the necrotic regions.

In the context of immune profiling, in our syngeneic model, we observed a significant induction of granzyme B, important for T cell receptor induced cell death, and inflammatory myeloperoxidase (MPO) in RFA treated tumors. Our comprehensive assessment of RFA side and non-RFA side contralateral tumor growth revealed RFA is a promising treatment for metastatic disease and primary tumors. Characterization of the TME revealed RFA treatment significantly changes the TME including induction of collagen and CD31+ cells indicating RFA directly or indirectly promotes angiogenesis, which may improve drug delivery. Our data on the non-RFA contralateral side tumors revealed similar changes in abundance of necrosis as well as changes in the TME. We quantified significant increases in collagen and CD31+ vessels on the contralateral side indicating Th1-mediated responses generate a potent alteration of the TME in non-RFA side contralateral tumors. To determine immune cell subsets in the non-RFA tumors, we used imaging mass cytometry (IMC) which revealed abundant anti-tumor macrophages were a significant component of the non-RFA side contralateral Th1 response. Histologic assessment of a resected tumor from a patient that received three rounds of RFA validated our murine preclinical findings. The ablation region had abundant collagen staining by trichrome and CD31+ vessels were adjacent to the RFA region. We also observed granzyme B and MPO staining in the parenchyma adjacent to the ablated pancreatic tissue.

To study changes in the TME, we used a proteome profiler array on lysates from control and RFA treated tumors. Consistent with elevated anti-tumor immunity, we observed a significant induction of C5/C5a, a prominent proinflammatory anaphylatoxin generated by cleavage of C5 during the compliment cascade ^32^. C5a promotes cytokine release and has been shown to promote endothelial cell chemotaxis and angiogenesis. Notably, C5a in non-small cell lung cancer has been shown to promote immunosuppression as blockade of C5a receptor decreased ARG1, CTLA-4, LAG3 and PDL1 in lung cancer models ^33^. As we observed increased C5/C5a in our RFA responsive tumors, these data suggest while acute anti-tumor local and systemic responses to RFA occur early after RFA, C5a may exert immunosuppressive mechanisms and impair complete response long term after RFA and highlight the need for pre-clinical evaluation of combination chemotherapy and immunotherapy with RFA to achieve improved responses in patients.

In addition to C5a, IL-23 was significantly increased in non-RFA contralateral tumors compared to control KPC tumors. IL-23 producing macrophages promote immunity to bacteria and exert anti-microbial functions ^34,35^. These data are consistent with our acute anti-tumor observation and IMC data showing a significant induction of F480+CD44+ macrophages which are known to produce IL-23. However, emerging data indicate elevated IL-23 promotes metastasis, growth of neoplastic epithelium, immunosuppression and suggest IL-23 is a target for cancer therapy ^36^. While correlative, directly related to our observations, IL-23 elevates matrix metalloprotease MMP9 and has been shown to elevate angiogenesis and has been shown to diminishes infiltration of CD8 T-cells ^37^. As IL-23 was significantly increased in RFA treated tumors, combined targeting of IL-23 with RFA may improve overall responses in patients with PDAC.

In addition to the proteome profiler, we used RPPA to evaluate proteomic changes in the non-RFA tumors compared to controls. Cdc42, GATA6, IGFRb and p-Akt^T308^, all drivers of PDAC growth and survival, were significantly decreased in responding non-RFA/contralateral tumors compared to controls. Surprisingly, a number of proteins known to have tumor suppressor function were significantly increased in non-RFA tumors including DDR1, INPP4b, MIG6 and EphaA2 ^38–41^. Notably, Drp1, a mitochondrial fusion GTPase that is required for Ras driven glycolysis and PDAC growth, was also significantly increased in contralateral tumors indicating tumor intrinsic mechanisms are elevated that need to be targeted for improved responses with RFA ^42^.

One observation was an increased FC tumor growth rate in non-responsive RFA treated tumors. The observed elevated FC tumor growth rates are consistent with previous reports showing incomplete RFA can increase tumor growth rates through significant induction of myeloid suppressor cells, increased new metastasis and sustained local inflammation in patients with colorectal cancer liver metastasis ^43^.

In this study, we wanted to evaluate RFA responses in a highly relevant preclinical model of PDAC. We have recently shown EUS-RFA is a promising therapeutic approach in combination with standard of care chemotherapy for patients with locally-advanced PDAC ^19^. We have now generated a preclinical ablation model which can be further evaluated for preclinical chemotherapy or immunotherapy combinations. In our current human clinical trial (www.clinicaltrials.gov NCT04990609), patients received three RFA sessions, which may generate sustained Th1 responses and reduce secondary systemic immunosuppression. In addition, we acknowledge and highlight other limitations of our preclinical data as these mice only received one RFA treatment and were not treated with chemotherapy or immunotherapy. The observed significant induction in CD31+ vasculature and angiogenesis suggest RFA may be a novel treatment to increase drug delivery to both local RFA treated tumors and distant metastasis. Future preclinical experiments evaluating RFA responses using multiple RFA treatments or in combination with IL-23 inhibitors, anti-PD1/PD-L1 or CD40 agonists will determine if these combination strategies promote survival for PDAC patients, especially patients with locally advanced or metastatic disease where innovative therapeutic options are needed to improve survival.

## Supporting information

Faraoni_Supplemental Figures

## ABBREVIATIONS

RFA: Radiofrequency ablation
TME: Tumor microenvironment
EUS: Endoscopic ultrasound
CAF: Cancer associated fibroblasts

