## Supplementary material for "Radiofrequency ablation remodels the tumor microenvironment and promotes systemic immunomodulation in pancreatic cancer": Faraoni_Supplemental Figures

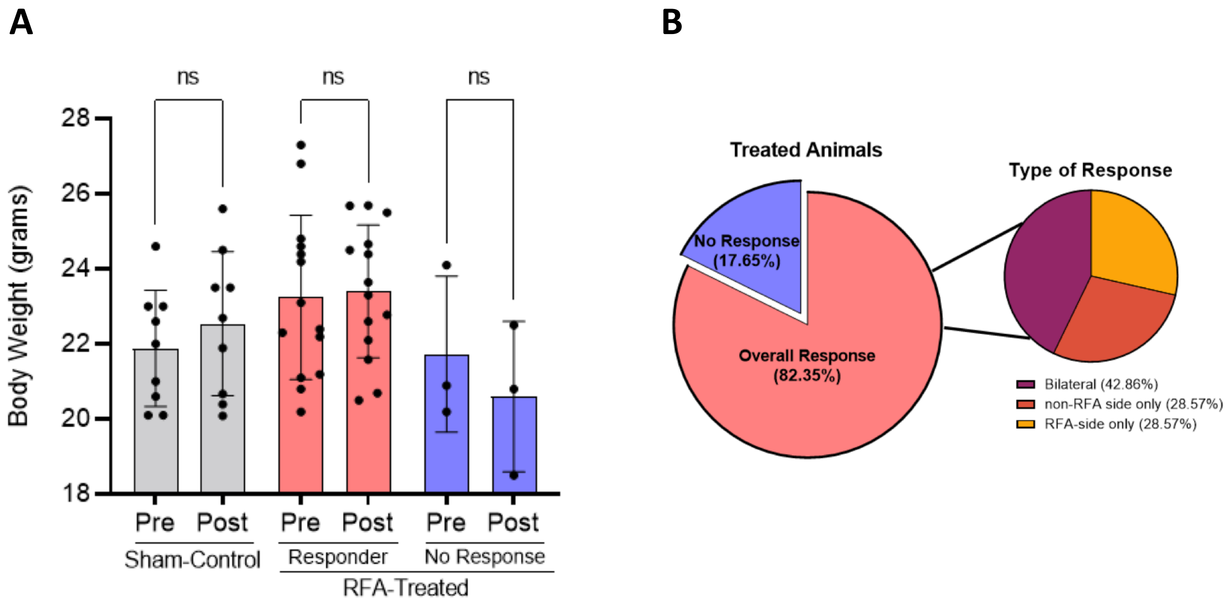


**Supplementary Figure 1. (A)** Body weight. Body weight was monitored before and after treatment. No differences were observed between control (*n*=8), responder (*n*=14) and no response (*n*=3) mice. Data was analyzed by paired 2-way ANOVA. Interaction ($P<0.05$). **(B)** Radiofrequency ablation treatment response *in vivo*. *See details in Table 1*.

**Supplementary Figure 2**

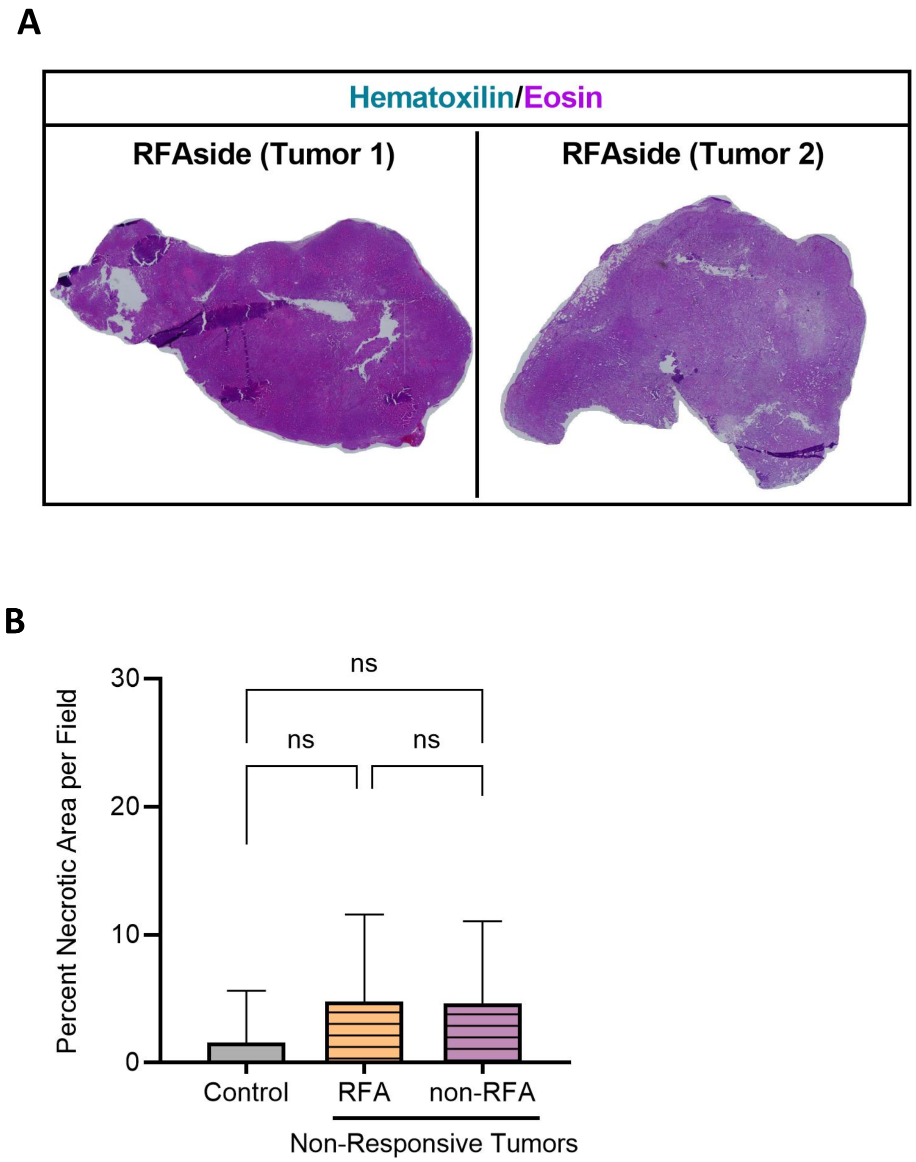


**Supplementary Figure 2. (A) Necrosis in non-Responding tumors from responding mice. A)** No response mice. Representative composite H&E staining of RFA side tumors from non-Responder mice. Necrosis quantification did not show significance in any of the samples analyzed (data not shown) suggesting technical failure in RFA procedure in these no response mice. **B)** ImageJ quantification of necrosis in KPC control (*n=8*), RFA (*n*=2) and non-RFA (*n=2*) non-Responding tumors. No differences were found in any of the groups. Data was analyzed by One-way ANOVA to determine statistical significance.

**A**

### RPPA Data

*Significantly downregulated in non-RFA contralateral tumors*

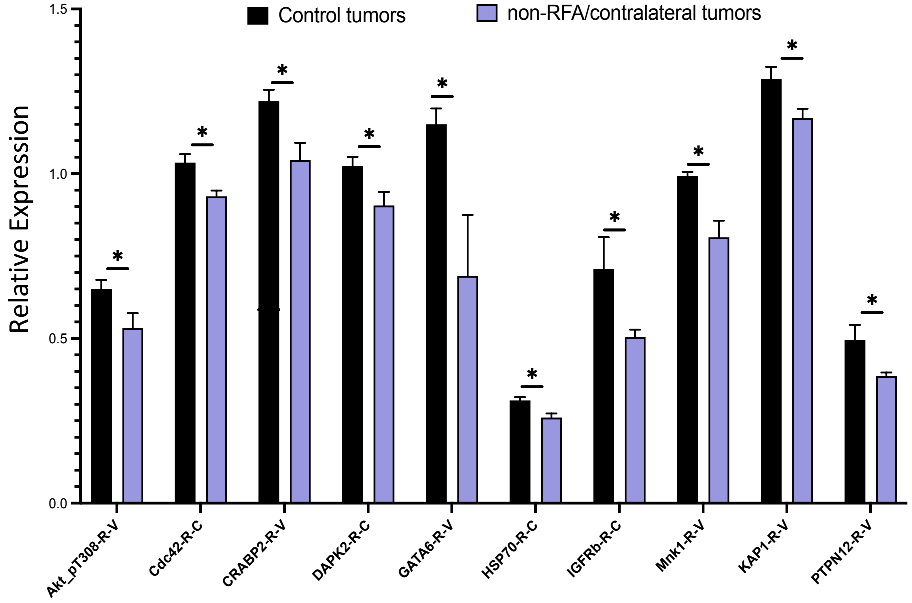


**B**

*Significantly upregulated in non-RFA contralateral tumors*

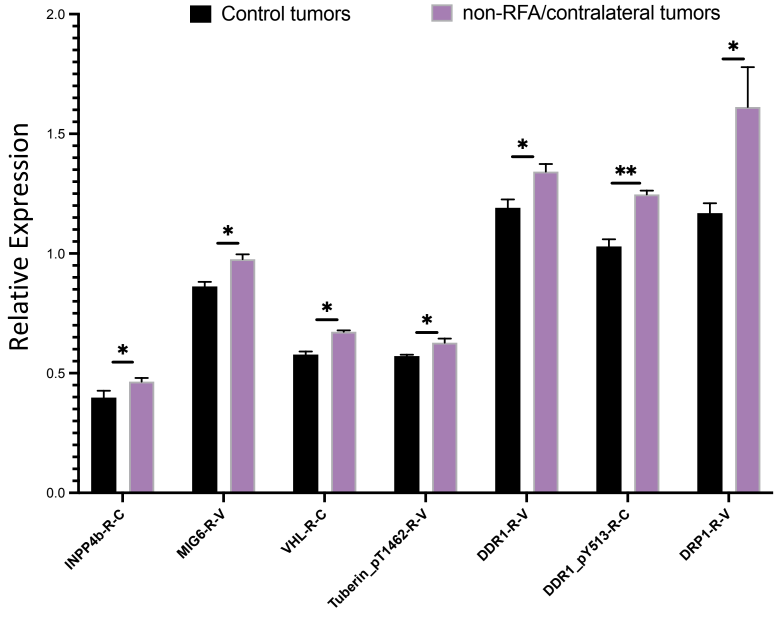


**Supplementary Figure 3. RPPA identifies pathways altered in contralateral tumors compared to control tumors. (A)** Proteins significantly downregulated include pAkt$^{T308}$, Cdc42, CRABP2, DAPK2, GATA6, HSP70, IFGRb, Mnk1, KAP1 and PTPN12 were all significantly decreased in non-RFA tumors (*p<0.05*). **(B)** INPP4b, MIG6, VHL, pTuberin$^{T1462}$, pDDR1$^{Y513}$ and DRP1 are significantly increased in non-RFA tumors compared to control tumors (*p<0.05*). Unpaired student's t-tests were used for comparisons.

| Antibody | Dilution IHC | Species | Catalog Number | Source | Antigen Retrieval |
|---|---|---|---|---|---|
| Granzyme B (mouse) | 1:250 | Rat | 13-8822-82 | Thermo Fischer | Vector Laboratories, H-3300 |
| Cleaved Caspase 3 | 1:400 | Rabbit | 9661 | Cell Signaling | Vector Laboratories, H-3300 |
| MPO (human and mouse) | 1:100 | Rabbit | 208670 | Abcam | Vector Laboratories, H-3300 |
| CD31 (human and mouse) | 1:200 | Rabbit | 281583 | Abcam | Abcam 100X Tris-EDTA pH 9 |
| NIMPR-14 | 1:200 | Rat | 2557 | Abcam | Vector Laboratories, H-3300 |
| Granzyme B (human) | 1:150 | Rabbit | 255598 | Abcam | Abcam 100X Tris-EDTA pH 9 |

**Supplementary Table 1. Antibodies used for IHC.**
